# First occurrence of Ranavirus in Scandinavian peninsula

**DOI:** 10.1101/2025.01.17.633642

**Authors:** B Thumsová, N Chondrelli, AE Valdés, R Eghbal, J Höglund, A Laurila, J Bosch, M Cortázar-Chinarro

## Abstract

Emerging infectious diseases (EIDs) pose a major threat to global amphibian populations, contributing to widespread mortality and species extinctions. Among EIDs, those caused by fungal infection pathogens *Batrachochytrium dendrobatidis* (*Bd*) and *B. salamandrivorans* (*Bsal*), and viral infections of the genus Ranavirus (*Rv*), represent the most significant threats to amphibian biodiversity. Here, we test for occurrence of *Rv* infection in three different locations in southeastern Sweden. Using a quantitative PCR (qPCR) assay, complemented by a secondary PCR-based validation method targeting the viral major capsid protein gene (*MCP*) and additional five partial sequences, we detected *Rv* infection in two of three locations and in five out of 43 individuals tested. This is the first confirmed record of *Rv* occurrence reported at such high latitudes in north Europe and the first one from Scandinavian peninsula, contributing important insights into infection prevalence in northern amphibian populations. These findings establish a basis for further research for the conservation of these vulnerable populations.

## Introduction

Emerging infectious diseases (EIDs) are a major cause of amphibian mortality worldwide and are implicated in the extinction of several amphibian species (Skerratt et al. 2007, Cunningham et al. 2017). Infectious diseases, particularly the fungal infection by *Batrachochytrium dendrobatidis* (*Bd*) and *B. salamandrivorans* (*Bsal)* and viral infection by *Ranavirus* (*Rv*), are considered ones of the most significant threats to global amphibian biodiversity (Skerratt et al. 2007, Duffus et al. 2015). The growing number of reports of *Rv* epizootics has recently raised concerns on account of their considerable population-level impacts (Price 2015, Bosch et al. 2021). Both pathogens are temperarature-dependent and, therefore, climate is one of the key environmental factors influencing their emergence, including infections caused by *Rv*, within *Iridoviridae* family (Epstein 2001, Cohen et al. 2017, Price et al. 2019, Thumsová et al. 2022a).

*Ranavirus* have been reported in several ectotherm taxa, including fish (Costa & Holmes 2024), amphibians (Davis et al. 2019, Thumsová et al. 2024), and reptiles (Brunner et al. 2015, Stöhr et al. 2015, Wirth & Ariel 2020), both in wild populations and in captivity. They were first isolated from Northern leopard frogs (*Lithobates pipiens*) in the USA in the 1960s (Granoff et al. 1965). In the late 1970s, the first occurrence of *Rv* was reported in European amphibians (Mişcalencu et al. 1981, Fijan et al. 1991). Since then, the detections of *Rv* has continued to rise both across Europe and globally (reviewed by [Price et al. 2017, Campbell et al. 2020]) with their presence now confirmed on all continents inhabited by amphibians (Marschang et al. 2024). Despite the global distribution, their origin of these viruses remains unknown to date (Thumsová et al. 2022).

Currently, the International Committee on Taxonomy of Viruses (ICTV) recognizes seven species of *Rv* circulating globally and infecting different taxa (Waltzek et al. 2024). In amphibians, these viral species are classified into three main groups, namely the *Frog Virus 3* (FV3-*like*) group, the most well-characterized group within the family *Iridoviridae*, alongside the *Common Midwife Toad Virus* (CMTV-like) and *Ambystoma Tigrinum virus (*ATV-*like*) ranaviruses. CMTV-*like* viruses impact a diverse range of species and life stages (Campbell et al. 2020) and appear to be the primary cause of *Rv*-induced mortality and morbidity in southern and central Europe (Price et al. 2017, Thumsová et al. 2022). CMTV-*like* viruses were first identified during investigations of a major mortality event in Picos de Europa National Park, Spain, in 2005, marking their initial detection in mainland Europe (Price 2014). Despite reported mass mortalities caused by *Rv*, Price et al. (2017) suggest that CMTV-*like Rv* are likely underreported in many regions across Europe. FV3-*like* viruses have been reported in amphibian species across the northern Iberian Peninsula and the United Kingdom (Price et al. 2017). Several studies have shown that CMTV-*like* and FV3-*like* viruses can co-occur, with a significant chance of recombination between them, potentially resulting in hybrid viruses that may be more severe than either CMTV-*like* or FV3-*like* viruses (Price et al. 2014, Price 2015, Claytor et al. 2017, Rosa et al. 2017). The most northern occurrence or *Rv* was documented in Danish edible frogs (*Pelophylax kl. esculentus*) in 2009 (Ariel et al. 2009). And, more recently, in a juvenile moor frog (*Rana arvalis*) in Latvia in 2022 (Di Marzio et al. 2024). Outside Europe, northern detections of *Rv* have been reported in northeastern Canada (Aoust-Messier et al. 2015) and more recently in Western Siberia in Russia (Lisachov et al. 2022).

Generally, populations residing at northern latitudes display less genetic diversity as compared to those from former refugial areas (Hewitt 1999). Previous studies suggest that amphibian species inhabiting northern latitudes may exhibit increased susceptibility to disease, potentially due to reduced overall genetic variation in these populations (Cortázar-Chinarro et al. 2017). In support of this, Meurling et al. (2024) found that *Bd* infection had a greater impact on common toad *Bufo bufo* individuals in northern populations, with higher observed mortality rates than in southern localities. Here, we investigated the presence of *Rv* in three high-latitude amphibian populations in the province of Kalmar (southeastern Sweden) to provide a foundational basis for understanding *Rv* occurrence and offering an opportunity to design targeted studies to investigate its distribution in northern amphibian populations.

## Methods

### Sample collection

On 23-27 April 2022, we collected a total of 86 samples (43 skin swabs and 43 tissue samples from toe clips) from 43 breeding *B. bufo* adults across three populations in the province of Kalmar (n=15 in Stjärnamo [population A; X=56.757863, Y= 16.06023], n=15 in Kindbäcksmåla [B; X= 56.507112, Y= 15.754273], and n=13 in Påryd [C; X= 56.608585, Y= 15.952671]; Table 1, Figure 1). We used a new pair of gloves for each toad collected and placed them in individual containers until skin and tissue sampling. Each toad was swabbed for infection detection with five strokes across each ventral and dorsal surface, front and hind limbs and hind toes. After swabbing, toads were rinsed twice with 140ml MilliQ water and had a piece of webbing clipped from their hind toes. Tissue samples were preserved in 96% ethanol and stored in freezers at -20°C. Swabs were kept cold during sampling and subsequently transferred to a -80°C freezer for long-term storage until samples were processed in the lab.

**Table 1.**
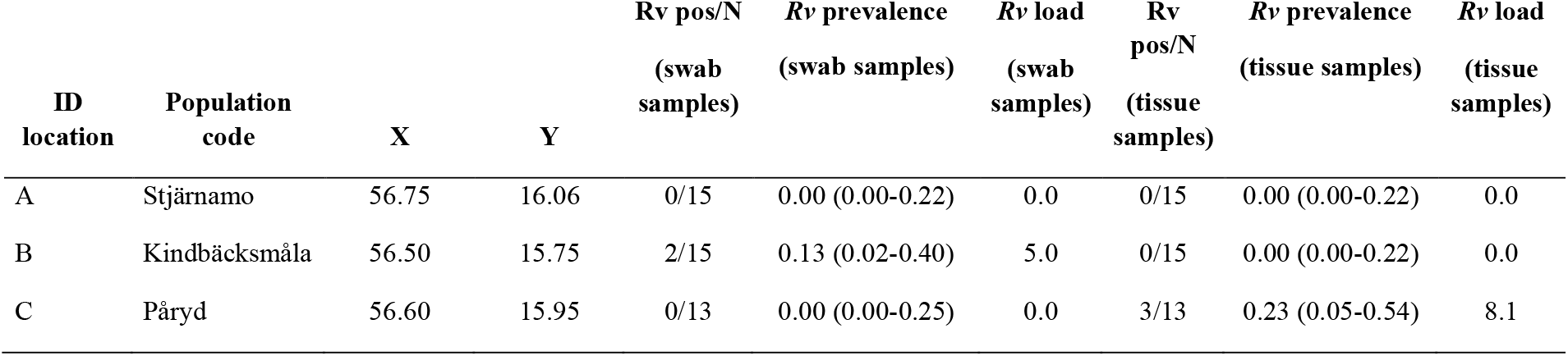
Geographic coordinates of Bufo bufo populations for Ranavirus (Rv), along with infection prevalence (with Clopper-Pearson confidence intervals in parentheses) and mean infection intensity expressed in genomic equivalents (GE) of virions.

| ID location | Population code | X | Y | Rv pos/N<br>(swab samples) | Rv prevalence<br>(swab samples) | Rv load<br>(swab samples) | Rv pos/N<br>(tissue samples) | Rv prevalence<br>(tissue samples) | Rv load<br>(tissue samples) |
| --- | --- | --- | --- | --- | --- | --- | --- | --- | --- |
| A | Stjärnamo | 56.75 | 16.06 | 0/15 | 0.00 (0.00-0.22) | 0.0 | 0/15 | 0.00 (0.00-0.22) | 0.0 |
| B | Kindbäcksmåla | 56.50 | 15.75 | 2/15 | 0.13 (0.02-0.40) | 5.0 | 0/15 | 0.00 (0.00-0.22) | 0.0 |
| C | Påryd | 56.60 | 15.95 | 0/13 | 0.00 (0.00-0.25) | 0.0 | 3/13 | 0.23 (0.05-0.54) | 8.1 |

**Figure 1.**
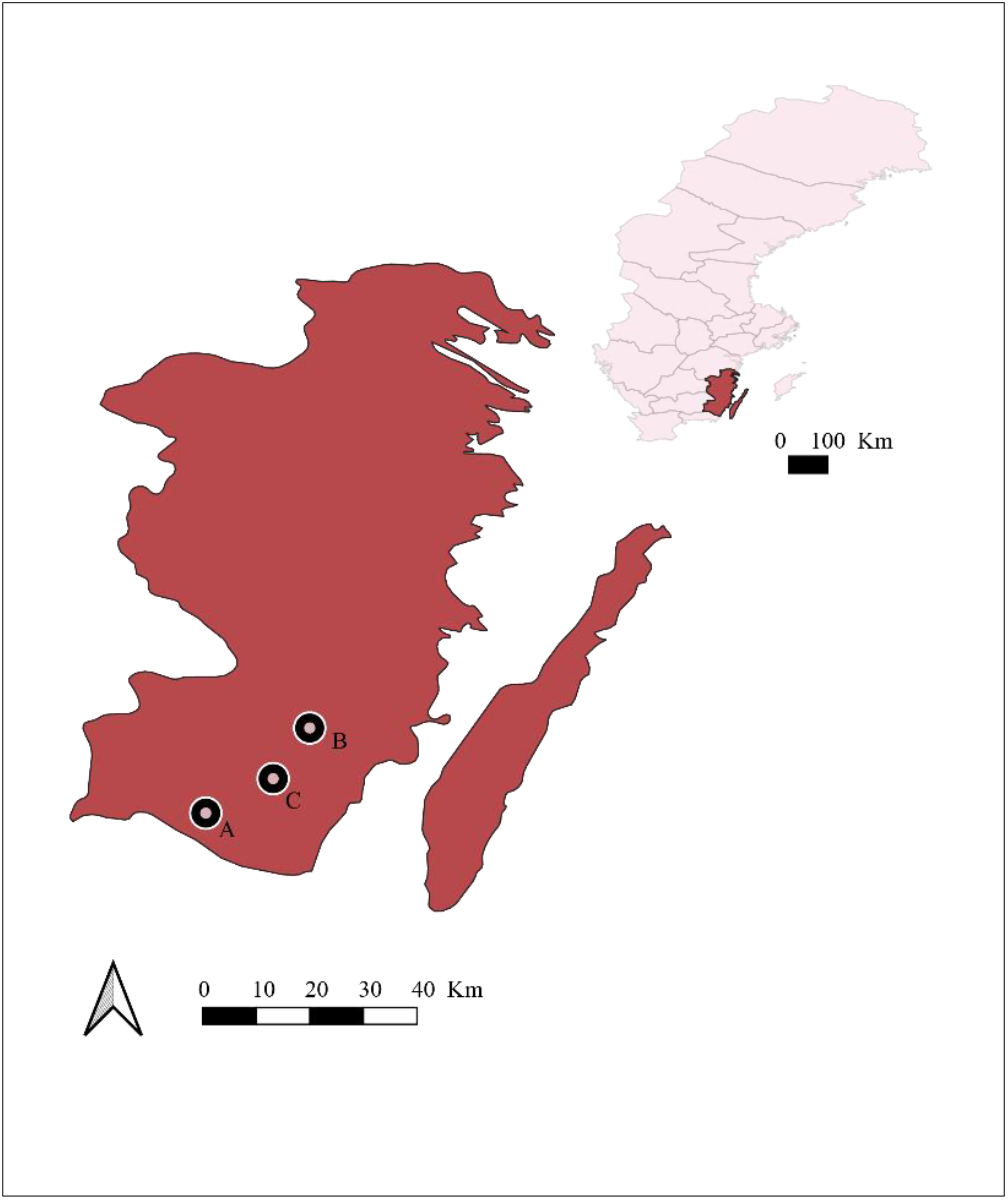
Map illustrating the sampling locations: (A) Stjärnamo, B (Kindbäcksmåla) and (C) Påryd in the southwest province of Kalma (Sweden).

### DNA extraction and infection detection

We extracted DNA from toe clips of each specimen using the DNeasy Blood and Tissue Kit (Qiagen GmbH, Hilden, Germany) following the manufacturer’s protocol. Skin swabs were processed for DNA extraction using the PowerSoil Pro Kit (Qiagen GmbH, Hilden, Germany), in accordance with the manufacturer’s instructions. For the detection and quantification of *Rv*, we used Real-time TaqMan PCR assays on a MyGo Pro machine, following the protocol by Leung et al. (2017), which targets a 97 base pair region of the viral Major Capsid Protein (*MCP*) gene. We ran all samples against negative and positive controls with known concentrations of genomic equivalents (GE) of virions from 0.1 to 10,000,000 in log10 increments. We considered a sample positive when the infection load was equal to or higher than the lower positive control, and the amplification curves presented robust sigmoidal shapes. We calculated 95% Clopper-Pearson (hereafter referred to as 95% CI) confidence intervals for site using Epitools (Brown et al. 2001).

For a double validation of the method, we subjected all positive tested samples to further PCR reactions to amplify the viral *MCP* gene using primers 4 and 5 from (Mao et al. 1997) as well as partial coding sequences from *Rv* open reading frames (ORFs) 22L (GenBank accession number AFA44926), 58L (AFA44964), 59R (AFA44965), 82L (AFA44988), and a region covering a noncoding sequence and the start of 13R (AFA44917), using primer sequences available in Rosa et al., (2017). We ran reactions in individual 0.2 mL tubes in 15 uL total volumes comprising 7.5 uL Platinum™ II Hot-Start PCR Master Mix 2X (Invitrogen), 0.75 uL of 10 mM stock of each primer, 4 uL of nuclease-free water, and 2 mL of template DNA. Following the methods used in Price et al. (2014) and Thumsová et al. (2022), we set the annealing temperature several degrees below the calculated annealing temperature for the primers. So, we ran the samples on the GeneAmp PCR System 9700 (Thermofisher Scientific Inc.) with the following settings for the viral *MCP* gene: an initial denaturation at 95°C for 10 min, followed by 35 cycles of denaturation at 95°C for 45 s, annealing at 52ºC or for 45 s and extension at 72°C for 45 s, with a final 7 min extension step at 72°C before holding at 4°C. To amplify the five additional loci (ORFs 13R, 22L, 58L, 59R, and 82L), we set the annealing temperature to 60°C. When weak or faint bands were observed during amplification, the number of PCR cycles was increased to 45 to enhance product yield. We electrophoresed PCR products on a 2 % agarose gel stained with GelRed (Biotium). We purified PCR amplicons with E.Z.N.A Gel extraction kit (Omega-Biotek) that there further submitted to Macrogen Spain for Sanger sequencing. The retrieved DNA sequences were blasted against the NCBI Reference Genomes Database (RefSeq_genomes; ref_viruses_rep_genomes) using the BLASTn program with the blastn algorithm (Camacho et al. 2009).

## Results

### Infection detection, load and prevalence

We detected *Rv* infection in 5 out of 43 sampled individuals (prevalence = 0.12, 95% CI: 0.04−0.25) using either swabs or toe clip tissue samples (Table 1). Specifically, 2 of 15 individuals from Population B (prevalence = 0.13, 95% CI: 0.02−0.40) and 3 out of 13 tissue samples from Population C (prevalence = 0.23, 95% CI: 0.05−0.54) tested positive. The mean infection load was 5.0 GE in positive individuals sampled by swabs and 8.1 GE in those sampled by tissue. None of the infected individuals showed visible signs of disease at the time of sampling.

### PCR amplifications of MCP gene

The five positive samples (40,41,43,61,74) were subjected to and additional PCR reaction to amplify the *MCP* gene and partial sequences of *Rv* genome. Although some amplification resulted in weak or multiple bands, for all samples, a positive band corresponding to *Rv* was detected in at least one of the targeted regions (See Table 2). Due to the low infection load found in our samples, sequences produced low-quality reads with noisy chromatograms, resulting inconsistent sequences that failed to align with known references in BLASTn. However, BLAST results for the *MCP* gene in samples 40 and 61 showed sequence similarities to CMTV-like, FV3-like, Bohle iridovirus (BIV), ATV and other ranaviruses.

**Table 2.** Result of polymerase chain reaction (PCR) of the viral major capsid protein gene (MCP, 16L) and the partial sequences from *Ranavirus* (*Rv)* open reading frames (ORFs): 22L (GenBank accession number AFA44926), 58L (AFA44964), 59R (AFA44965), 82L (AFA44988), and a region covering a noncoding sequence and the start of 13R (AFA44917) using primer sequences available in Rosa et al. (2017). Yes indicates when a positive band corresponding to (*Rv*) was detected in at least one of the targeted regions.

| Target gene | Locus<br>(ORF ref.) | Sample ID |  |  |  |  |
| --- | --- | --- | --- | --- | --- | --- |
|  |  | 40 | 41 | 43 | 61 | 74 |
| Major capsid protein gene | 16L | yes | yes | yes | yes | no |
| Proliferating cell nuclear antigen<br>gene | 22L | no | no | no | no | no |
| Hypothetical protein gene | 58L | yes | yes | yes | yes | yes |
| Hypothetical protein gene | 59R | no | no | no | yes | no |
| p31k gene | 82L | no | no | no | no | no |
| Hypothetical protein gene | 13R | no | no | no | yes | no |

## Discussion and Future directions

We investigated *Rv* occurrence in three locations within the Kalmar province in southern Sweden, an area previously monitored for *Bd* (Cortazar-Chinarro et al., unpubl.). These locations were chosen as part of ongoing projects in which two cases of amphibian mass mortality were observed in spring 2024, although no mass mortality events have been recorded in the area for *B. bufo* in Southern Sweden. However, this area is recognized as a regional hotspot for amphibian diversity in Sweden, representing the northernmost distribution range for several amphibian species primarily distributed in central Europe but with limited ranges in Scandinavia such *bufotes viridis* (Sillero et al. 2014). Therefore, the emergence of any new pathogen in this area could compromise the survival of many of these species in the near future.

We detected *Rv* infection at low intensities in five individuals across two of the three sampled locations, using two different approaches: qPCR and PCR amplification of the *MCP* gene, along with additional partial sequences from *Rv* genome. Additionally, BLAST analysis of the *MCP* gene identified several ranaviruses in two of the five positive samples, one from each of the two infected populations.These findings confirm and validate the presence of *Rv* in at least these locations for the first time in Sweden (Marschang et al. 2024). Field studies conducted on wild populations and laboratory-based experimental research have demonstrated that *Rv* prevalence and infection intensity are directly influenced by temperature fluctuations and other exogenous environmental factors (Miller et al. 2011, Price et al. 2019, Thumsová et al. 2022). For instance, general patterns of disease spread shown that *Rv* is negatively associated to canopy cover. A recent study conducted in the Brazilian rainforest revealed unconventional patterns in *Rv* disease dynamics, showing a positive correlation between forest cover and *Rv* infection prevalence. The study observed that tadpoles had higher *Rv* infection loads in areas with greater vegetation cover (Ruggeri et al. 2024), which could be potentially linked to the immune responses of different host species to infection [e.g., (Hoverman et al. 2011)]. Additionally, *Rv* has been found to be associated with high temperatures or sudden temperature increases in montane areas of the Northern Hemisphere [e.g., (Bosch et al. 2020, Thumsová et al. 2022b)]. Therefore, we suggest that the low infection intensity observed in this study may be attributed to the relatively mild environmental conditions during the sampling season in April, expecting higher infection rates around summer. At this stage, it is not possible to determine whether the temperatures during the breeding season are optimal for *Rv* proliferation, as this might be dependent on the host’s immune system condition. To confirm this and to capture episodes of peak infection or detect potential mass mortalities in the area, sampling efforts should be conducted at multiple time points throughout the breeding season, with a particular focus on the hottest periods of the year. These efforts should involve collecting skin swabs, buccal swabs, and internal organs from individuals at various life stages (e.g., adults, tadpoles, and metamorphs) in order to assess whether the infection affects individuals across different stages of their life cycle.

Both CMTV-*like* and FV3-*like* viruses have been documented in Europe. CMTV-like viruses have been reported in Belgium, France, Germany, the Netherlands, Portugal, Spain, and the United Kingdom, whereas FV3 viruses have been identified only in Denmark, northern Spain, Portugal, the Netherlands, and the United Kingdom (Marschang et al. 2024, Price et al. 2017). The review by Rodrigues et al. (2024) also noted the presence of *Rv* in Sweden, citing the study by (Stöhr et al. 2013). However, in Stöhr et al. (2013) *Rv* was detected in several *P. kl. esculentus* individuals collected for laboratory experiments from different ponds in Europe. Stöhr et al. (2013) characterized the *Rv* strains and identified two distinct ranaviruses clustering within the *Phelophylax esculenta* virus (REV-*like*) clade. The authors clarified that, based on comparisons of samples collected before and after the individuals were transported to the laboratory, the *Rv* source population originated from Germany, not Sweden. Therefore, our study represents the first evidence for the presence of *Rv* in Sweden. Due to the low viral intensity detected in our samples sequencing produced poor-quality reads for most of the cases. Although, confirmed the presence of *Rv* in two cases, we cannot, at this stage, resolve which exact *Rv* group (CMTV-*like* or FV3-*like*) the detected virus belongs to.

Several other *Rv* species have also been reported across Europe, including the Epizootic Hematopoietic Necrosis Virus (EHNV) and the European North Atlantic *Rv (*ENARV). However, many detected viruses remain poorly characterized (Marschang et al. 2024). A significant challenge lies in screening for specific *Rv* species or strains in regions where multiple strains or species coexist, which may lead to under detection and introduce biases in interpreting the host and geographical range of individual *Rv* species. Given previous findings of *Rv* infection-related mortalities in northern latitudes and their conservation implications, we emphasize the need for a continuous amphibian disease monitoring system in these regions. Such a system should incorporate comprehensive metagenomic screening to enhance the quality and resolution of viral genomes, thereby increasing the chances of detecting a broader range of viral strains over time.

## Acknowledgments

Funding was provided by Helge Ax:on Johnsons Stiftelse and Stiftelsen för Zoologisk Forskning to NC and MC.

